# Semantic Surprise Predicts the N400 Brain Potential

**DOI:** 10.1101/2022.05.31.494099

**Authors:** Alma Lindborg, Lea Musiolek, Dirk Ostwald, Milena Rabovsky

**Author notes:** These authors contributed equally (Alma Lindborg, Lea Musiolek).

## Abstract

Language is central to human life; however, how our brains derive meaning from language is still not well understood. A commonly studied electrophysiological measure of on-line meaning related processing is the N400 component, the computational basis of which is still actively debated. Here, we test one of the recently proposed, computationally explicit hypotheses on the N400 – namely, that it reflects surprise with respect to a probabilistic representation of the semantic features of the current stimulus in a given context. We devise a Bayesian sequential learner model to derive trial-by-trial semantic surprise in a semantic oddball like roving paradigm experiment, where single nouns from different semantic categories are presented in sequences. Using experimental data from 40 subjects, we show that model-derived semantic surprise significantly predicts the N400 amplitude, substantially outperforming a non-probabilistic baseline model. Investigating the temporal signature of the effect, we find that the effect of semantic surprise on the EEG is restricted to the time window of the N400. Moreover, comparing the topography of the semantic surprise effect to a conventional ERP analysis of predicted vs. unpredicted words, we find that the semantic surprise closely replicates the N400 topography. Our results make a strong case for the role of probabilistic semantic representations in eliciting the N400, and in language comprehension in general.

**Significance Statement:** When we read or listen to a sentence, our brain continuously analyses its meaning and updates its understanding of it. The N400 brain potential, measured with electrophysiology, is modulated by on-line, meaning related processing. However, its computational underpinnings are still under debate. Inspired by studies of mismatch potentials in perception, here we test the hypothesis that the N400 indexes the surprise of a Bayesian observer of semantic features. We show that semantic surprise predicts the N400 amplitude to single nouns in an oddball like roving paradigm with nouns from different semantic categories. Moreover, the semantic surprise predicts the N400 to a much larger extent than a non-probabilistic baseline model. Our results thus yield further support to the Bayesian brain hypothesis.

## Introduction

Language conveys meaning, but how our brains achieve this mapping from words and sentences to meaning is not yet well understood. The N400 brain potential has been studied extensively as a measure of on-line meaning-related processing, using a multitude of meaning-related materials such as sentences, word pairs, single words, pictures, sounds, and even mathematical symbols (Kutas and Federmeier, 2011). However, despite very large amounts of data, the computational processes underlying N400 amplitudes are still actively debated. There are various verbally descriptive hypotheses on the underlying mechanism of the N400 such as that N400 amplitudes may reflect the difficulty of lexical access to a word in memory (Lau et al., 2008), the difficulty to integrate an incoming word into the sentence context (Baggio et al., 2008), or semantic inhibition (Debruille, 2007). In recent years there has been a growing interest in linking N400 amplitudes to computational models in order to resolve this debate.

It has been suggested that information processing can be analyzed and modeled at three levels: computational, algorithmic, and implementational (Marr, 2010). So far, models of the N400 have been mostly implemented as neural network models (Rabovsky and McRae, 2014; Frank et al., 2015; Brouwer et al., 2017; Rabovsky et al., 2018; Fitz and Chang, 2019), which operate (roughly) at the “algorithmic” level of analysis (sometimes considered “implementational”). Some of these neural network modeling studies have argued that the models’ N400 correlates (i.e., the measures obtained from the neural network models, which have been used to simulate N400 amplitudes) approximate constructs such as semantic prediction error and/or Bayesian surprise (Rabovsky and McRae, 2014; Rabovsky et al., 2018; Fitz and Chang, 2019), concepts that we jointly refer to as semantic surprise here. This idea has been taken up by verbal theories of the N400 (Bornkessel-Schlesewsky and Schlesewsky, 2019; Kuperberg, 2016) and seems well in line with recent studies using large scale deep learning language models, which highlight the crucial role of prediction for language in general (Schrimpf et al., 2021) and the N400 in particular (Heilbron et al., 2020; Lindborg and Rabovsky, 2021). However, the theory that N400 amplitudes reflect semantic surprise has not yet been tested with a model operating at Marr’s “computational” level, that explicitly implements probabilistic Bayesian computations.

On a conceptual level, linking the N400 to semantic surprise connects with advances in the study of the mismatch negativity (MMN), a negative EEG potential arising in response to oddball stimuli (Näätänen et al., 1978). Recent studies have suggested that the underlying mechanisms of the MMN could be explained by predictive coding (Garrido et al., 2009), with Bayesian surprise (Itti and Baldi, 2009) as a possible predictor of MMN amplitude (Ostwald et al., 2012). Similarly, amplitudes of the oddball sensitive P3 component have been linked to Bayesian surprise (Mars et al., 2008; Kolossa et al., 2015; Visalli et al., 2021; Modirshanechi et al., 2019) and predictive coding accounts suggest that ERP amplitudes generally reflect surprise at different levels of representation (Friston, 2005). Importantly, while ERPs in perceptual oddball paradigms have been explicitly modeled as Bayesian surprise; concerning the N400, the hypothesis that it reflects semantic surprise has not yet been tested with explicit probabilistic modelling. This is the goal of the current study.

Specifically, we devise a simple probabilistic model of semantic knowledge – a Bayesian sequential learner – which continuously tracks the probabilities of different semantic categories during the course of a word reading experiment and updates its knowledge when presented with new information. We test whether the model-derived semantic surprise, technically defined here as Bayesian surprise with regard to the model’s internal semantic representation, can predict trial-by-trial N400 amplitude in human subjects taking part in an oddball like roving paradigm experiment with words from different semantic categories. We further investigate whether the temporal and spatial EEG signature of the semantic surprise are consistent with the N400 obtained from standard ERP analyses.

## Methods and Materials

### Participants

Forty subjects (36 female) with a mean age of 24 (SD = 3.71) participated in the experiment. All participants were native German speakers with normal or corrected-to-normal vision and were right-handed according to 12 items of the Edinburgh Handedness Questionnaire (Oldfield, 1971). The study was approved by the Ethics Committee of Freie Universität Berlin’s Department of Psychology and all participants provided written informed consent before participation.

### Stimuli

The stimuli consisted of 100 German nouns falling into 10 semantic categories, with 10 nouns from each category. The semantic categories were trees, vegetables, birds, land animals, landscapes, furniture, transportation, tools, kitchen utensils and clothes. For the purpose of the experiment we constructed the 10 discrete categories by selecting nouns which maximised the semantic similarity of words within a given category using the semantic features from the GermaNet lexical-semantic net (Hamp & Feldweg, 1997; Henrich & Hinrichs, 2010). The stimuli were restricted to have no more than one commonly used meaning and to be reasonably well-known. It was verified in separate one-way factorial ANOVAs that the categories did not significantly differ on any of the following control variables: absolute type frequency (*F* (9) = 1.37, *p* = 0.12) and number of orthographic neighbours according to Coltheart’s (*F* (9) = 1.34, *p* = 0.23) and Levenshtein’s (*F* (9) = 0.90, *p* = 0.53) definitions – all extracted from dlexDB German language corpus (Heister et al., 2011) – as well as the number of letters (*F* (9) = 1.30, *p* = 0.25).

In order to make sure that participants were paying attention to the stimuli, they were required to perform a control task consisting of detecting non-words. These non-words had no orthographic neighbours in German. A full list of the word and non-word stimuli can be found in the Supplementary material.

### Experimental design

The stimuli were presented in a roving paradigm, in which a sequence of nouns belonging to one semantic category (e.g., different trees) were followed by a sequence of nouns from a different category (e.g., tools), and so on. With this paradigm, we can regard the contrast of “standard” and “deviant” trials akin to that of an oddball paradigm, where the first stimulus in a sequence is a “deviant” (unpredicted) stimulus and the last stimulus in a sequence is a “standard” (predicted). In total, 500 category sequences of varying length (min = 4, max = 8, mean = 6 nouns) were constructed such that the mean sequence length was balanced across categories, and the number of repetitions (30) and probability of occurring at a given position in a sequence was balanced across word items. The sequences were presented in a randomised order, yielding a total of 3000 stimulus presentations during the experimental session.

Additionally, 200 catch trials consisting of non-words were presented during an experimental session. Subjects were tasked with pressing a designated key whenever a non-word appeared on the screen, in order to make sure they were paying attention to the stimuli. The non-words were randomly interspersed between the nouns but restricted to occur after 8-24 words and never directly before or after a standard (last in sequence) stimulus. Subjects were asked to press a given key whenever a non-word occurred in the stimulus presentation. The laterality of the key press was balanced across participants and subjects’ performance in the non-word task was continuously monitored during the experiment.

Each word or non-word was presented at the centre of the screen for one second, followed by a fixation cross during a random, uniformly distributed time interval between 650 and 950 ms, resulting in an average ISI of 800 ms.

### Procedure

Participants were seated in front of a computer screen on which the stimuli were presented at a visual angle of at least 2.3 degrees, delivered using the Matlab Psychophysics 3 Toolbox (Brainard, 1997). Before the start of the experiment, participants completed 20 practice trials to make sure they understood the task. The experiment was divided into 12 blocks, each comprising 250 words. After each block the participants’ performance in the non-word task was displayed on the screen to increase their motivation. Participants could take breaks between the blocks if they wished.

### EEG recording and pre-processing

EEG data were recorded at a sampling rate of 2046 Hz using a 64-channel active BioSemi system with a channel layout corresponding to the extended 10/20 system, with an additional two mastoid and four ocular channels. Pre-processing was performed in Brain Vision Analyzer 2 (Brain Products GmbH, Gilching, Germany). First, noisy channels were identified in six participants (1-2 channels per participant) and were interpolated from the surrounding channels using spline interpolation. Second, the continuous data were re-referenced to the mastoid average, intervals of no activity were automatically detected and marked for removal, and the data of each channel was band-pass filtered using a second order zero-phase Butterworth IIR filter with a low cut-off frequency of 0.023 Hz and a high cut-off frequency of 30Hz. The channel-wise data were then segmented into epochs starting 100 ms before stimulus onset and ending 900 ms after stimulus onset, and subsequently downsampled to 512 Hz. Automatic ocular artefact correction was performed on the segmented data using the Gratton & Coles method. Furthermore, trials containing values higher than 200 *µ*V or lower than − 200 *µ*V, gradient steps higher than 50 *µ*V/ms and/or a value difference of more than 200 *µ*V in a 200 ms interval were marked as artefactual and excluded from further analyses. In total, 0.95% of trials were marked for rejection during preprocessing. Trials were then baseline-corrected to the 100 ms interval preceding the stimulus onset. Upon artefact detection at conventional levels of data resolution, the data epochs were shortened to include the -100 ms to 800 ms interval relative to stimulus onset and downsampled to 128 Hz. This additional data compression was performed in order to facilitate the time-resolved trial-by-trial analysis.

For the N400 amplitude analysis, we averaged the signal over a fronto-central region of interest (ROI) comprising 15 electrodes (see Figure 2), corresponding to the sensors showing the largest difference between standard and deviant trials in the N400 time interval defined as 300-500 ms after stimulus onset. As the ROI we found was slightly more anterior compared to standard N400 experiments (eg. Hodapp and Rabovsky 2021; Rabovsky et al. 2012), we additionally conducted the same analyses on a more centro-parietal ROI, which yielded the same basic pattern of results of a much better prediction of N400 amplitudes by semantic surprise as compared to a non-Bayesian baseline model (please see Supplementary material).

Two main types of EEG analyses were conducted: a classical ERP analysis contrasting the standard and deviant conditions, and a trial-by-trial analysis using our model of semantic surprise to predict the single-trial EEG.

### ERP analyses

In the ERP analyses, we first averaged the signal spatially over the ROI. Subsequently, the standard (last item in sequence) and deviant (first item in sequence) trials were averaged separately by subject (500 trials per subject and condition before artifact rejection). Finally, for the statistical testing we estimated the average N400 amplitude by subject and condition by averaging the signal over the time window 300-500 ms after stimulus onset. We hypothesised that deviant trials would have a more negative N400 than standards, which was tested by subjecting standard and deviant trials to a two-tailed paired sample t-test, at a significance level of *α* = 0.05.

### Trial-by-trial analyses

#### Semantic surprise model

We derived our measure of trial-by-trial semantic surprise from a Bayesian sequential learner model based on a categorical representation of the stimuli, in which each word falls into one of 10 discrete categories. The model simulates the behaviour of an experiment participant who, at each trial, compares the observed word to their current beliefs about the semantic context and subsequently adjusts its beliefs according to the new observation.

Following earlier work on the N400 (Rabovsky et al., 2018), we implemented our semantic surprise measure as the Bayesian surprise, which indexes the extent to which a stimulus causes the learner to update its beliefs (Modirshanechi et al., 2021). If, for example, a learner has previously been presented with only land animals, seeing another land animal will not cause a large adjustment to the learner’s beliefs, resulting in a small BS. Seeing a tool will, however, prompt the learner to reconsider the probabilities of the respective semantic categories, and will thus cause a larger Bayesian surprise. The BS is defined as the Kullback-Leibler (KL) divergence between the model’s beliefs prior to, and after, the presentation of a stimulus (Modirshanechi et al., 2021). An example of how BS varies over a sequence of stimuli in the roving paradigm is displayed in the upper panel of Figure 1.

**Figure 1.**
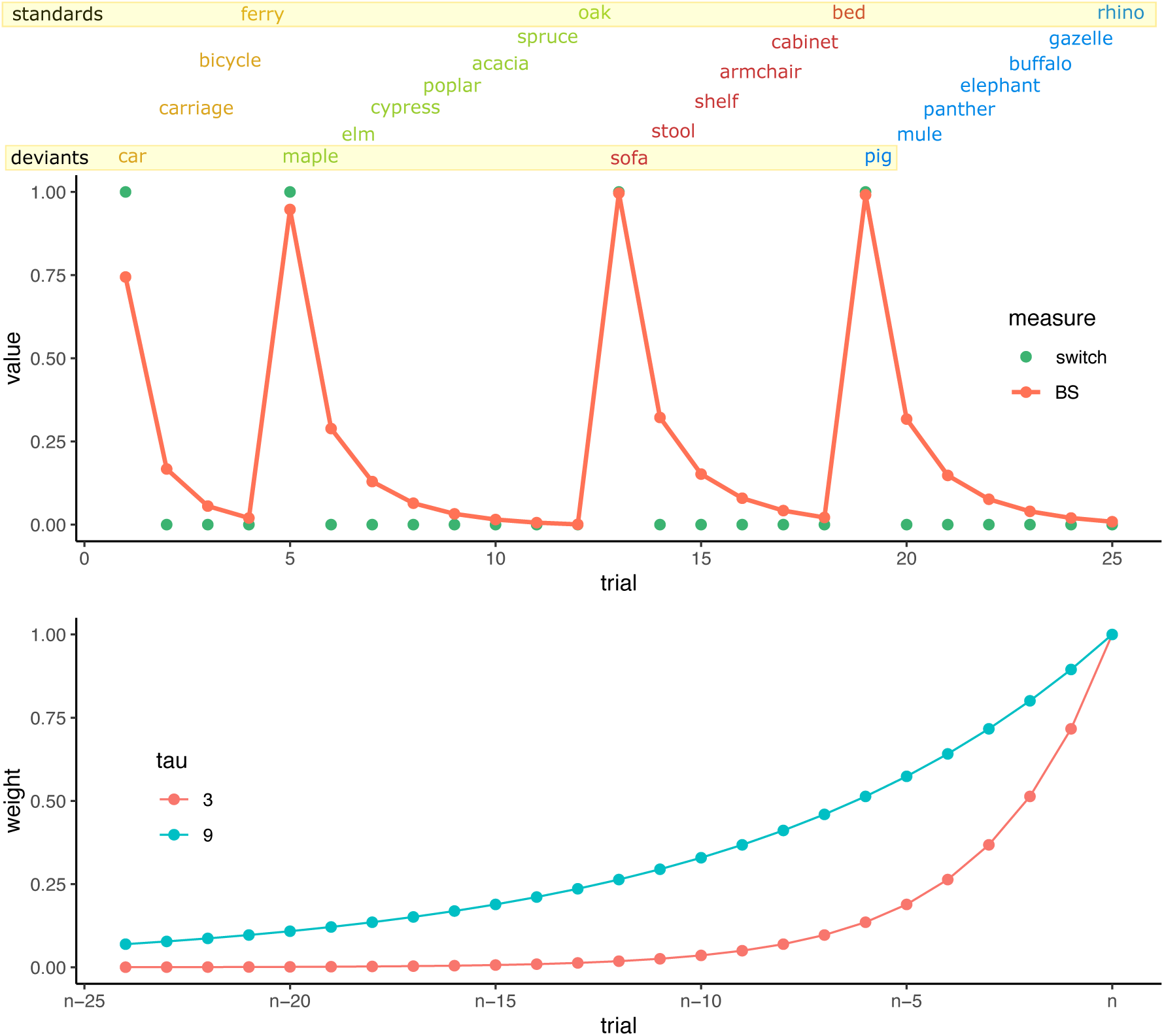
Experiment structure and semantic surprise. Upper panel: nouns from a given category are presented in sequences of varying length, and the Bayesian learner model adjusts its beliefs after each word. The first word in each sequence is relatively unexpected and gives a large semantic surprise – these trials are labelled deviants for the ERP analyses. The semantic surprise decreases gradually throughout the sequence. The last trial in each sequence is labelled as standards for the ERP analyses. Lower panel: the memory weighting function for different values of tau. A weight of 1 means the trial is fully remembered, whereas a weight of 0 means it is fully forgotten. With a lower value of *τ*, only relatively recent trials are remembered, whereas a larger *τ* allows for more trials to be remembered.

Formally, we implemented our sequential learner as a Dirichlet-Categorical model (Gijsen et al., 2021), in which the likelihood of observing a stimulus of category *i* given the current beliefs follows a categorical distribution:

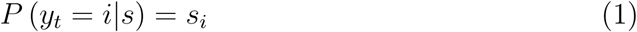

Here, the current beliefs 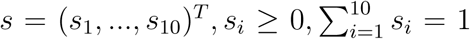 are the probabilistic representation of category probabilities for the ten semantic categories used in the experiment. We assume that the learner encodes its uncertainty about the category membership of a given stimulus using a Dirichlet distribution

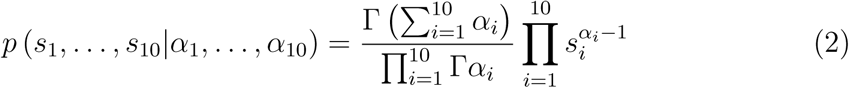

with the parameters *α*_*i*_, …, *α*_10_. After observing stimulus *y*_*t*_, we apply Bayes’ rule to find the posterior estimate of the beliefs:

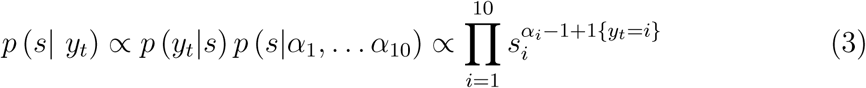

Since the Dirichlet and categorical distributions are conjugate, the posterior distribution is also a Dirichlet distribution, with the *α*_*i*_ parameters updated to incorporate the new observation. Thus, given an initial value 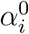 (fixed to 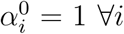 in our experiment) we can compute the current beliefs of the model at any time point *t* as

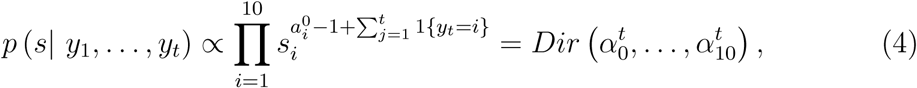

where

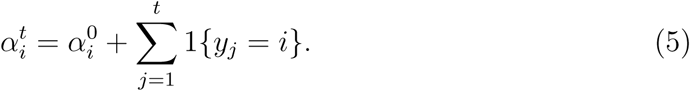

However, as this calculation of 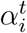 assumes infinite memory of previous trials, and thus doesn’t reflect the biological constraints of memory, we apply an exponential memory decay function to simulate the forgetting process. Applying exponential memory decay to the trials, we get

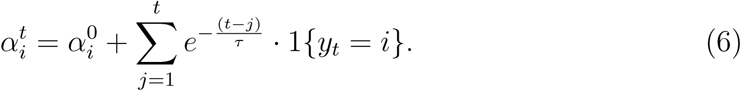

The memory decay function is visualized for different values of *τ* in Panel B of Figure 1. The Bayesian Surprise at trial *t* can now be directly calculated as the KL divergence between the distribution of the model’s beliefs prior and posterior to observation of stimulus *y*_*t*_:

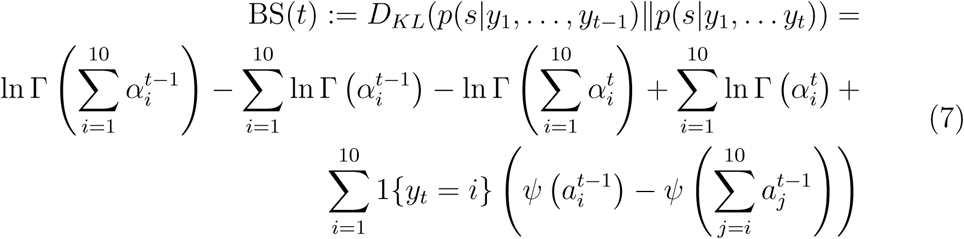

with *ψ* referring to the digamma function and Γ the gamma function.

As a non-probabilistic baseline model, we use a category switch measure, which assumes maximal semantic surprise when the current word comes from a different category compared to the previous one, and no semantic surprise otherwise. The category switch measure is plotted with green dots for the example sequence in the upper panel of Figure 1.

Semantic surprise was calculated separately on each subject’s stimulus sequence, and min-max scaling was applied so that the surprise always fell on the [0, 1] interval. The value of *τ* was estimated from the EEG data of all subjects simultaneously.

#### N400 amplitude analysis

In the single trial analysis, the N400 amplitude from each trial was entered into the statistical analyses, excluding trials with an absolute N400 amplitude exceeding 75*µ*V (0.19% of trials) which were considered outliers, leaving 117432 trials for analysis. We tested whether the N400 amplitude significantly depends on semantic surprise, using a mixed linear model with the N400 as dependent variable, the semantic surprise as predictor and random intercepts for subject and lexical item. Formally, the statistical model was thus

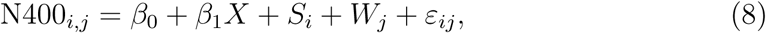

where the N400 of subject *i* and word *j* depends on the intercept *β*_0_ and the regression coefficient *β*_1_ times the semantic surprise *X*, plus the random intercepts *S*_*i*_ and *W*_*j*_ for subject *i* and word *j*, respectively. The random intercepts are assumed to be drawn from centered normal distributions: 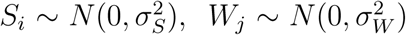 and *ε*_*ij*_ ∼ *N*(0, *σ*^2^).

Note that the model-derived semantic surprise depends on the forgetting parameter *τ*, as shown in Equation (6) and illustrated in the lower panel of Figure 1. We determined the value of *τ* by maximising the fit of the linear mixed model in Equation (8). This was done by computing the semantic surprise at each integer value between *τ* = 1 and *τ* = 15, fitting a linear mixed model of the N400 to the semantic surprise at each value of *τ* and selecting the value that provided the best prediction of the N400 amplitude as indicated by the F-statistic of the semantic surprise predictor. Thus, *τ* was optimised over the whole dataset simultaneously, yielding a single value of *τ* for all subjects. This value was subsequently used for the temporally and spatially resolved analyses.

#### Time-resolved trial-by-trial analysis

In the time-resolved single trial analysis, we wanted to test to what extent semantic surprise could predict single-trial EEG in a time resolved manner, an approach which is commonly called encoding models (Di Liberto et al., 2015; Smith and Kutas, 2015; Holdgraf et al., 2017). This approach entails fitting a regression model where each time sample of the EEG is predicted by the semantic surprise. However, fitting a mixed effects model such as that defined in Equation (8) to each time point independently would result in a prohibitively large number of free parameters to estimate. Given 40 subjects, 100 words and 114 time points, the number of random intercept parameters would amount to (40 + 100) · 114 = 15690, in addition to a fixed intercept and slope parameter for each time point (114 · 2 = 228), yielding a total of 15918 parameters. Since such a high-dimensional model would be prone to significant over-fitting to the data, we instead adopted a regularised regression approach using ridge regression, which constrains the model in order to decrease its variance (Bishop, 2006). In ridge regression, the parameters 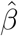 are estimated as

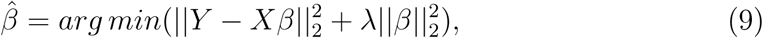

where the first term minimising the difference between *Y* (the observations at each time point) and *X* (the design matrix) times *β* (containing an intercept and slope term for each time point) is the ordinary least squares minimisation problem and the second term penalises solutions with a large *ℓ*2 norm. The shrinkage parameter *λ* determines the amount of constraint applied to the regression. We computed a separate ridge regression model on each subject’s data, thus letting 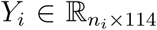 contain the *n*_*i*_ ≤ 3000 trials of 114 time points from subject *i*, 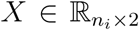 be the design matrix containing a column of ones for the intercept and a column containing the model semantic surprise at each trial, and lastly letting *β*_*i*_ ∈ ℝ_2×114_ be the matrix of regression intercepts and slopes for each time point. This reduced the problem to the estimation of a total of 40 · 114 · 2 = 9120 free parameters. See Heilbron et al. 2020; Di Liberto et al. 2015; Smith and Kutas 2015 for similar regression-based approaches to single-trial EEG modelling.

In order to constrain each subject’s regression parameters equally, we determined *λ* by fitting a single regression model on the data from all participants for each value of *λ* on the log-spaced interval [10^−10^, 10^−9^, …, 10^10^] and picking the value yielding the lowest generalisation error, as estimated by leave-one-out cross-validation over trials using the implementation from Scikit-learn (Pedregosa et al., 2011). Subsequently, subject-specific encoding models were computed with the pre-determined shrinkage parameter.

Encoding coefficients, i.e. the slope parameters estimated for each subject, were tested statistically for each time point by a two-tailed one-sample t-test (or a Wilcoxon signed-rank test, if the sample did not pass D’Agostino and Pearson’s normality test) with a significance level of *α* = 0.05, corrected for multiple comparisons using the Bonferroni-Holm method (see eg. Heilbron et al. 2020 for a similar approach). We visualised the topography of the effect by similarly computing a linear encoding model predicting the mean voltage per channel in the 300-500 ms time window. All trials from all subjects were used for this ridge regression, with the shrinkage parameter again determined by leave-one-out cross-validation.

### Code and Data Accessibility

All code and (summarised) data necessary for producing the presented results will be made publicly available upon publication. The raw EEG data are available upon request.

## Results

### ERP analysis

The results of the condition-based analysis are shown in the upper panels of Figure 2. In our region of interest, we find that the ERP to deviant trials significantly differs from that of standards in the pre-selected time window 300-500 ms after stimulus onset. In line with our hypothesis, the N400 was significantly larger for deviant trials compared to standard trials (*t*(39) = −3.08, *p* = 0.004). The topography of the effect is plotted in Panel B, displaying the difference between the deviant and standard conditions in the 300-500 ms time window. The sensors included in the ROI are marked in red.

**Figure 2.**
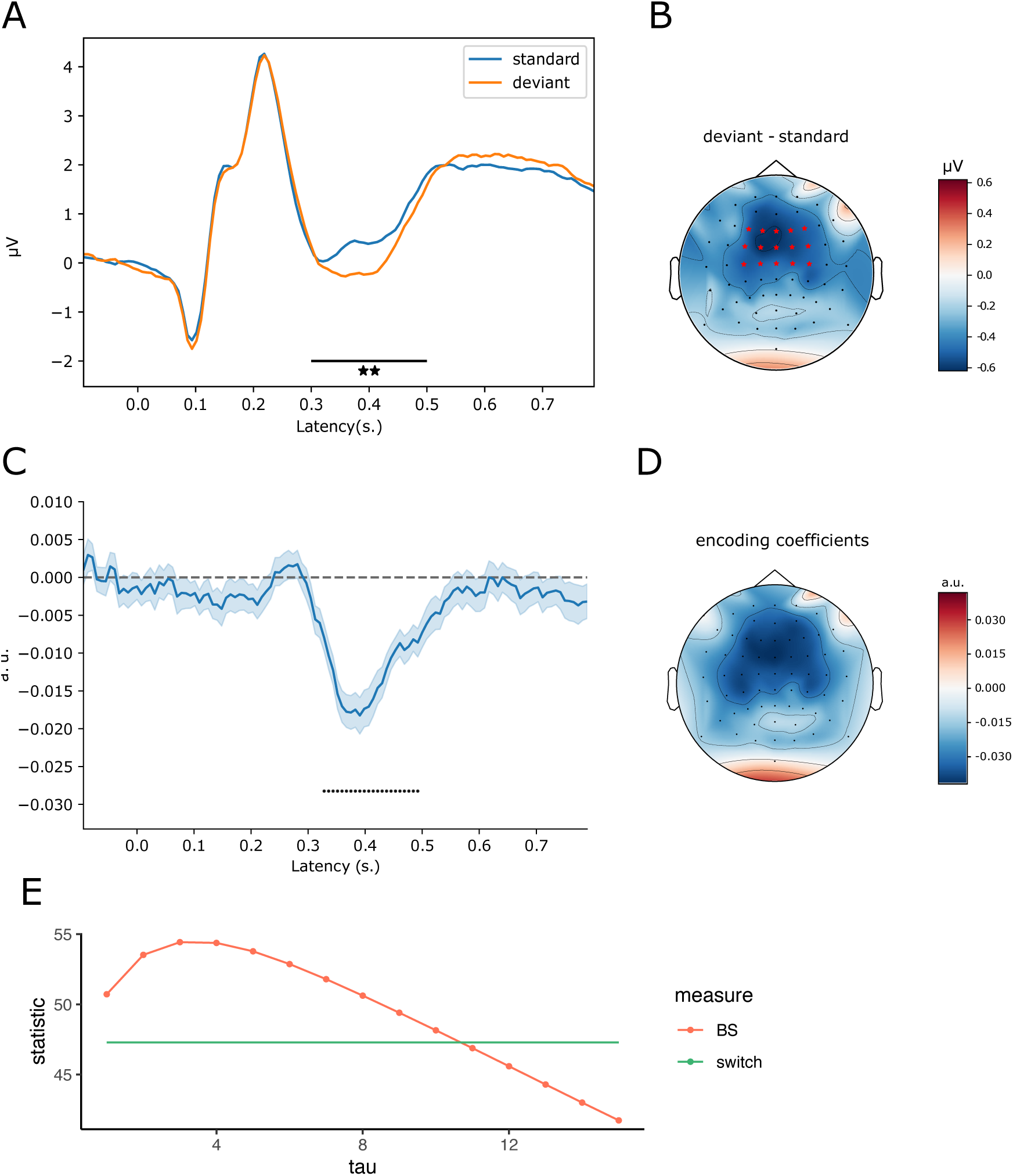
Top row: ERP analysis. Panel A shows grand-average ERP for standard and deviant trials over the ROI, with the 300-500 ms time window used in the statistical test marked with a black line. Panel B shows the grand average difference between deviant and standard trials 300-500 ms after stimulus onset. The sensors belonging to the ROI are marked in red. Middle and bottom rows: single-trial analysis. In Panel C, the mean of the subject-specific encoding coefficients from the trial-by-trial analysis are shown for the ROI. Error bands represent the standard error of the mean. Time points where the coefficients significantly differ from zero are marked with dots. Panel D: topographic plot of encoding coefficients estimated on all subjects and trials in the N400 time window. Panel E: Trial-by-trial average N400 analysis. F-statistic as function of surprise measure and forgetting parameter.

### Trial-by-trial analysis

The mixed linear model analyses of the single trial N400 revealed that both the Bayesian surprise and the category switch measure predicted the N400 amplitude. A separate model was estimated for each regressor (one for category switch and one for Bayesian surprise at each value of *τ*) and an F-test of each model confirmed that the fixed effects were significant (*F*(1) *>* 41, *p <* 10^−9^ computed with Satterthwaite’s method). The results are plotted in Panel E of Figure 2. Applying model selection based on the Akaike Information Criterion (AIC) values (see eg. Anderson 2008 for details) of the best-fitting Bayesian surprise model (*τ* = 3) and category switch, we found that Bayesian surprise captured 97.25% of the combined explained variance of both models, whereas category switch only captured 2.75%. Thus, semantic surprise derived from the sequential Bayesian learner model predicts the N400 better than the non-Bayesian category switch model.

In the time-resolved single trial analysis, we used semantic surprise with the best fitting memory decay value (*τ* = 3) to predict the time continuous EEG of each participant, using ridge regression. The shrinkage parameter was set to *λ* = 1000 after optimisation on the complete data set and statistics were performed on subject-specific encoding models. The results are displayed in Panel C of Figure 2. After correction for multiple comparisons, the coefficients significantly differed from zero at 22 time points, all of which fell within the 300-500 ms time window and had negative coefficients. The topography of the effect, computed in a channel-wise encoding model of the N400 time window estimated on the full data set, is shown in Panel D of Figure 2.

## Discussion

In the current study, we have investigated the computational underpinnings of the N400 brain potential in language comprehension. We have tested whether a probabilistic model tracking the probabilities of semantic categories can simulate the N400 in a meaningful way. We have found that the Bayesian surprise, i.e. the amount of adjustment to the model’s internal semantic representation caused by observing a stimulus, significantly predicts the N400 amplitude in single trials. Moreover, investigating the temporal and spatial signature of the effect of semantic surprise, we have shown that it is specific to the time window and topography of the typical N400 effect, that is the difference between standard and deviant trials in the ERP analyses.

Our results are in line with models and theories suggesting that N400 amplitudes reflect semantic surprise (Rabovsky and McRae, 2014; Rabovsky et al., 2018; Lopopolo and Rabovsky, 2021; Kuperberg, 2016; Bornkessel-Schlesewsky and Schlesewsky, 2019). However, as noted in the introduction, so far these proposals were either based on verbally descriptive theories (Kuperberg, 2016; Bornkessel-Schlesewsky and Schlesewsky, 2019) or on neural network models operating at Marr’s algorithmic (partly considered implementational) level of analysis (Rabovsky and McRae, 2014; Rabovsky et al., 2018; Lopopolo and Rabovsky, 2021). Here, we complement the neural network based approach with an explicitly probabilistic Bayesian modeling approach operating at Marr’s computational level (see also Delaney-Busch et al. 2019, but Nieuwland 2021).

One advantage of this explicitly Bayesian approach is that it allows to address the issue to what extent predictions in language processing are probabilistic in the sense that they are reliant on estimates of uncertainty. This claim, which is closely related to the “Bayesian brain” hypothesis (Knill and Pouget, 2004), cannot be easily tested with current neural network-based language models, as even large, state-of-the-art neural network language models with near-human performance in language tasks (Vaswani et al., 2017; Radford et al., 2019) and interesting correpondences to the human language system (Frank et al., 2015; Schrimpf et al., 2021; Heilbron et al., 2020; Caucheteux and King, 2020; Michaelov and Bergen, 2020) lack an explicit representation of uncertainty. Though theoretical analogies exist between neural network models and Bayesian inference (McClelland, 2013), in practice, uncertainty quantification in neural network models is a non-trivial problem (Abdar et al., 2021). Our observer model, although simplistic in its feature space, demonstrates that the semantic surprise derived from a probabilistic observer model predicts the N400 far better than that of a non-probabilistic observer (only tracking the switches between categories).

The results of the current study set the N400 in relation to ERP components observed in perceptual oddball paradigms such as the MMN and P3, which have featured prominently in Bayesian accounts of brain function (Garrido et al., 2009; Ostwald et al., 2012). This relation suggests that similar probabilistic processing principles may apply across levels of representation, in line with the idea that ERPs might generally reflect surprise at different processing levels (Friston, 2005).

Although we would like to suggest that semantic surprise, indexing the on-line adjustments a person makes to their probabilistic representation of an utterance or text, is of general significance to understanding natural language, we have tested this proposition with a rather limited form of stimuli (nouns). The use of 10 discrete categories for our word stimuli and observer model is a clear limitation of the study. Clearly, this crude categorical representation of nouns does not correspond to the current theories of semantic categories, which frames categories as clusters in a high-dimensional space of semantic features rather than well-defined groups of objects (e.g., McRae et al. 1997). Our stimulus material was specifically selected to have high within-group and low between-group similarity, and was thus optimised for a categorical model. This experimental limitation enabled the use of a simple and computationally explicit Bayesian sequential learner model. However, a more general model could instead employ semantic feature vectors such as feature norms (McRae et al., 2005) or data-driven word embeddings. Such a design would also have the advantage of more closely approximating the continuous and graded (rather than discrete) grouping of concepts in natural language.

We have here used the term semantic surprise to denote the model’s Bayesian surprise; however, this is not the sole possible surprise measure. An alternative (though related) measure of semantic surprise briefly mentioned in the Introduction is the prediction error, which indexes the improbability of an event under the current beliefs. Thus, whereas the Bayesian surprise quantifies the discrepancy of an agent’s beliefs prior to and after stimulus presentation, the prediction error directly compares the stimulus to the prediction generated by the beliefs. The prediction error is defined as the negative log-likelihood of the stimulus under the current beliefs (Modirshanechi et al., 2021). We provide a detailed definition of prediction error as well as results of the single-trial analysis based on prediction error in the Supplementary Material.

Bayesian surprise and prediction error could not be disentangled by our model (post-hoc testing showed that the linear correlation between the measures is 0.96). We argue that this is not a general feature of the respective surprise measures, but rather that it is a consequence of our choice of experimental paradigm and computational model. Since the Bayesian surprise indexes the update of the model’s beliefs, it is highly dependent on the precision (inverse variance) of the probabilistic representation of the beliefs – the more certain the beliefs are, the smaller the update caused by an unexpected stimulus. The prediction error however is not affected by the precision of the beliefs in this way, since it only relies on the expectation value of the predictive distribution. Thus, future work might disentangle Bayesian surprise and prediction error, for example by varying the perceptual clarity or semantic ambiguity of the stimuli. Since a Bayesian learner updates its beliefs by Bayes’ rule, the extent to which evidence alters its beliefs depends on the relative precision of the evidence compared to the beliefs (cf. Gelman 2014). Therefore, an unpredicted stimulus (i.e. with large prediction error) which is not well perceived (such as a word presented in noise) or is semantically ambiguous (such as “penguin”) would cause a smaller update to the model’s beliefs compared to an equally unpredicted stimulus which is perceptually and semantically clearer (such as “robin” presented without noise).

In summary, we have shown that semantic surprise, estimated by a Bayesian sequential learner model, predicts the amplitude as well as topography of the trial-by-trial N400 brain potential. Moreover, we have demonstrated an approach to explicitly model the probabilistic processing which the N400 has long been claimed to indicate. We believe that extensions of this approach could further disentangle the complex computational processes supporting the human ability to seemingly effortless derive meaning from a conversation or a text.

## Supporting information

Supplementary Analyses

## Notes

### Competing Interest Statement

The authors have declared no competing interest.

