## Supplementary Analyses for "Semantic Surprise Predicts the N400 Brain Potential"

### Supplementary Material 1: Supplementary Analyses

May 31, 2022

#### Prediction error analyses

Here, we present results of single-trial EEG analyses using an alternate measure of semantic surprise, namely prediction error, using the sequential Bayesian learner model and the EEG data defined in the main article.

##### Definition of prediction error

Prediction error, or "puzzlement surprise" (Faraji et al., 2018), is defined as the negative log-likelihood of an observation $y_t$ given the current beliefs $s$. Under the Bayesian sequential learner scheme, $s$ is conditioned on the previous observations $y_1, \dots y_{t-1}$, and the predictive distribution is thus obtained by marginalising the over $s$:

$$P(y_t = i|\ y_1, \dots, y_{t-1}) = \int p(y_t = i|s)p(s|y_1, \dots, y_{t-1})ds = E[s_i|\ y_1, \dots, y_{t-1}], \quad (1)$$

which, given the Dirichlet distribution of $s$ over 10 categories, yields the prediction error:

$$PE(y_t) = -\ln \frac{\alpha_i}{\sum_{i=1}^{10} \alpha_i^{t-1}} \quad (2)$$

with $\alpha_i^{t-1} = \alpha_i^0 + \sum_{j=1}^{t-1} 1\{y_j = i\}$. Thus, in contrast to Bayesian surprise (BS), the prediction error (PE) only depends on the *relative* counts of previous observations, disregarding how strong the current beliefs are.

##### EEG analyses

Results of the EEG analyses are plotted in Figure 1. In the 300-500 ms time window and the ROI selected following ERP results, the mixed linear model analyses suggested that all measures (BS, PE and category switch) significantly predicted the N400 amplitude ($F > 41$, $p < 10^{-9}$ in each separate ANOVA, computed with Satterthwaite's method). Moreover, the F-values indicated that BS predicted the N400 amplitude slightly better than PE, and that both probabilistic surprise measures produced a better fit than the baseline model (Panel C). Model comparison using Akaike's information criterion confirmed that the best fitting model for BS (with $\tau = 3$) captured 59.28% of the total variance, whereas PE (at $\tau = 6$) stood for 39.04% of the total variance and the category switch model only 1.68%. Thus, BS indeed seemed to fit slightly better to the data compared to PE; however, the difference was not substantial. Panels A and B show the time-resolved and spatially

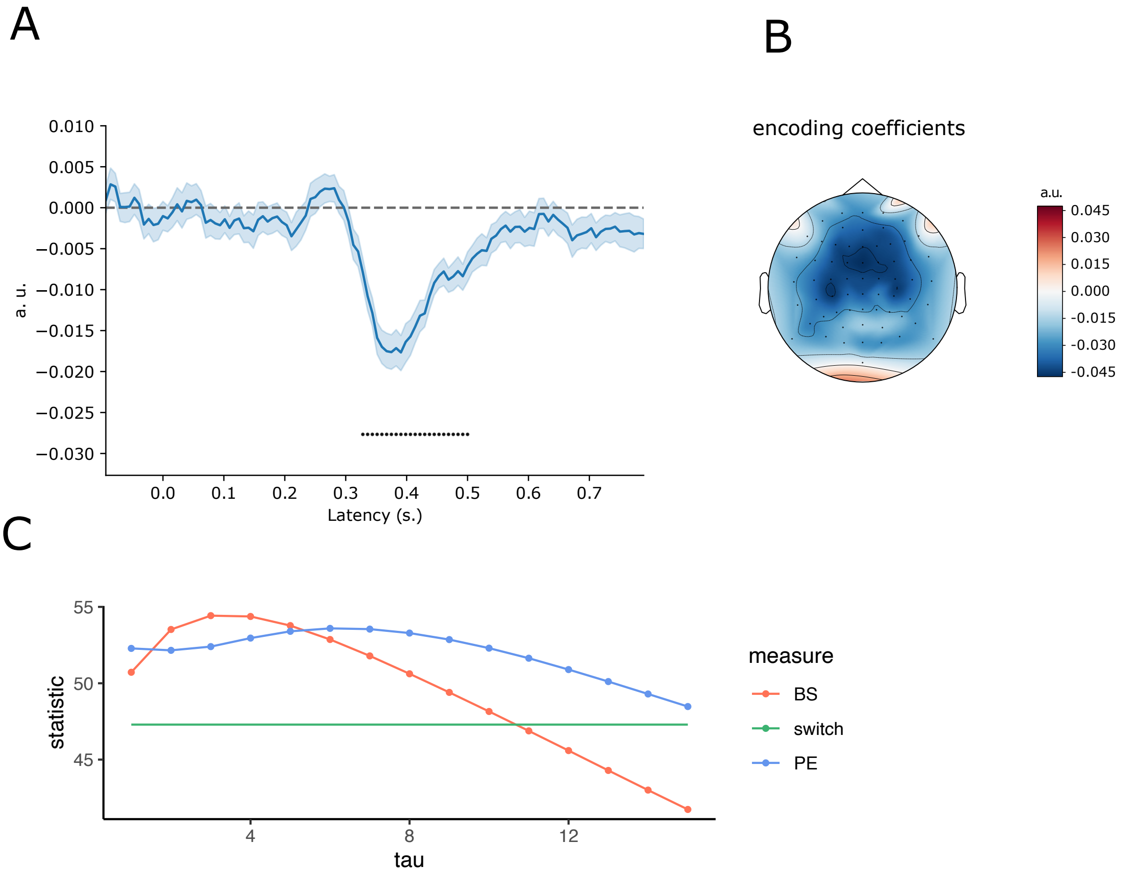


**Figure 1.** EEG analyses for the prediction error predictor. In Panel A, the mean of the subject-specific encoding coefficients are shown for the ROI. Error bands represent the standard error of the mean. Time points where the coefficients significantly differ from zero are marked with dots. Panel B: topographic plot of encoding coefficients estimated on all subjects and trials in the N400 time window. Panel C: Single trial average N400 analysis. ANOVA F-statistic for prediction error (PE), Bayesian surprise (BS) and the non-probabilistic baseline model (switch) as function of surprise measure and forgetting parameter.

resolved linear encoder analyses, both using PE as a predictor. Very much in line with the results for BS, the PE predicts a negative potential at 23 significant time points restricted to the $300 - 500$ ms time window. Further testing revealed almost perfect correlation between the measures; for $\tau = 6$ the correlation between BS and PE was 0.96, whereas for $\tau = 3$ the correlation reached 0.98.

#### Alternative ROI

The ROI presented in the main text was selected to maximise the difference between the standard and deviant ERPs. Although results on the topography of the N400 to single nouns are rather mixed (Ganis et al., 1996; Stróżak et al., 2016), this ROI is decidedly more anterior than a "standard" N400 ROI. Therefore, we additionally conducted the same statistical analyses for a more centrally located ROI comprising the 5 central electrodes on the FC, C and CP rows.

##### Single trial analyses

As in the main text, we fit a linear mixed model to the mean N400 amplitude over the ROI with the model surprise (Bayesian or non-probabilistic) as predictor and random intercepts for each subject and lexical item. Applying an F-test to the fixed effect in each model, we found that every regressor significantly predicted the N400 amplitude ($F > 40, p < 10^{-9}$). Picking the value of $\tau$ which maximises the fit for each probabilistic measure ($\tau = 8$ for BS and $\tau = 10$ for PE), we compared the model fits by their AIC values. This analyses indicated that the category switch model captured only 0.04% of the pooled model variance, whereas prediction error accounted for 95.39% and Bayesian surprise for 4.56%. Thus, in the data-driven ROI and central ROI alike, the probabilistic measures clearly provided a superior fit to the N400 compared to the non-probabilistic baseline. However, in contrast to the data-driven ROI, the prediction error predicted the N400 over the central ROI better than the Bayesian surprise.
